# Minimalistic mycoplasmas harbor different functional toxin-antitoxin systems

**DOI:** 10.1101/2021.01.21.427616

**Authors:** Virginia Hill, Hatice Akarsu, Rubén Sánchez Barbarroja, Valentina Cippà, Martin Heller, Laurent Falquet, Manfred Heller, Michael H. Stoffel, Fabien Labroussaa, Joerg Jores

## Abstract

Mycoplasmas are minute bacteria controlled by very small genomes ranging from 0.6 to 1.4 Mbp. They lack a cell wall and have been suggested to have progressed through reductive evolution from phylogenetically closely related Clostridia. They are known to colonize the respiratory tract or the urogenital tract among other organs and can cause chronic and subclinical diseases associated with long persistence of the causative agent. Toxin-antitoxin systems (TAS) are genetic elements that have been described for several respiratory and urogenital pathogens as well as for Clostridia, but never for pathogenic mycoplasmas. Here we describe for the first-time different types of TAS in a *Mycoplasma* pathogen, namely *M. mycoides* subsp. *capri*. We identified candidate TAS *in silico* via TASmania database. Two candidate TAS identified *in silico* and another candidate TAS suggested in a minimal cell based on transposon mutagenesis were systematically tested for their functionality in hosts with different phylogenetic distance using heterologous expression. Phylogenetic distance of the host used for heterologous expression influenced the outcome of the functional testing. We corroborated functionality of the three candidate TAS in *Mycoplasma capricolum* subsp. *capricolum*. Moreover, we confirmed transcription and translation of molecules of the TAS investigated during *in vitro* growth. We sequence analyzed 15 genomes of *M. mycoides* subsp. *capri* and revealed an unequal distribution of the TAS studied pointing towards dynamic gain and loss of TAS within the species.

**Author summary:** Mycoplasmas have a minimal genome and have never been shown to possess TAS. In this work we showed the presence of different functional TAS systems in *Mycoplasma mycoides* subsp. *capri*, a caprine pathogen for the first time. Sequence analysis of a number of *Mycoplasma mycoides* subsp. *capri* strains revealed a plasticity of the genome with respect to TAS carriage. This work paves the way to investigate the biological role of TAS (e.g. persistence, stress tolerance) during infection using mycoplasmas as a simple model organism. Since most mycoplasmas lack classical virulence factors such as exotoxins and go into a kind of stealth mode to evade the immune system, TAS are likely to contribute to the parasitic lifestyle of mycoplasmas and should be investigated in that respect. The availability of synthetic genomics tools to modify a range of *Mycoplasma* pathogens and well-established challenge models for the latter mycoplasmas will foster future research on TAS in mycoplasmas.

## Introduction

Mycoplasmas are the smallest bacteria that can replicate in axenic media reported so far. Their cell size and genome size are minute compared to most other bacteria. The genomes range from 0.6 to 1.4 Mbp, a feature that attracted researchers to use mycoplasmas as model organisms for synthetic cells [1] and minimal cells [2]. The small genome size has been attributed to a reductive evolution of mycoplasmas, which are lacking a cell wall and are phylogenetically closely related to spore-forming GRAM-positive bacteria, the Clostridia [3]. A characteristic feature of the mycoplasmas is their parasitic lifestyle, which is reflected by their inability to synthesize essential building blocks of life such as amino acids and hitherto their dependence on a host to provide these building blocks to fuel the anabolic pathways.

Mycoplasmas colonize different niches of their hosts such as mucous membranes of the respiratory tract, the ear canal or the urogenital system. The genus *Mycoplasma* encompasses important human pathogens such as *M. genitalium* [4] and *M. pneumoniae* [5] as well as pathogens of utmost veterinary importance such as *M. mycoides* subsp. *mycoides* [6], *M. capricolum* subsp. *capripneumoniae* [7] and *M. hyopneumoniae* [8]. Although a number of mycoplasmas cause diseases with a relatively short incubation time and high lethality as known for contagious caprine pleuropneumonia [9], many *Mycoplasma* infections are chronic and associated with a long persistence of the causative agent [10, 11].

Toxin-antitoxin systems (TAS) are chromosomal or extrachromosomal genetic elements, which can be grouped into six different types according to their mode of action [12]. All TAS encode a toxin (T), which is able to interfere with vital processes of the bacterial cell related to transcription, translation, replication and membrane integrity, as well as an antitoxin (A) that inhibits the activity of the toxin, so that it cannot interfere with the vital cellular processes. TAS have been described for a wide range of GRAM-positive and GRAM-negative bacteria including pathogens of the gastrointestinal and respiratory tract such as Clostridia and Mycobacteria, respectively. The role of TAS has been attributed to reduce metabolism during stress, hinder bacteriophages, stabilize genetic elements, and modulate the build-up of biofilms [13].

Pathogenic mycoplasmas are not known for classical exotoxins, the only exception is an ADP-ribosyltransferase (ART) activity-conferring toxin identified in *M. pneumoniae* [14]. So far, TAS have not been documented in mycoplasmas. The presence of a TAS in an engineered strain of *Mycoplasma mycoides* subsp. *capri* (*Mmc*) with a heavily reduced genome has been suggested based on polar effects of transposon insertions [15], but was never confirmed using heterologous expression or shown in its parental wild type strain GM12.

The goal of this study was to shed light on the presence of TAS in mycoplasmas, which are known for their minute genome size. Therefore, we chose *Mmc* GM12, a caprine pathogen [16] used to study virulence traits [17-19] and used as model organism for synthetic genomics. We performed an *in silico* analysis to identify candidate TAS, which were subsequently systematically tested for their functionality in hosts with different phylogenetic distance using heterologous expression. Thereafter, we examined the presence of the candidate TAS at the species and genus level and noticed an unequal distribution pointing towards horizontal gene transfer to be involved in dissemination of the TAS over different mycoplasmas.

## Results

### Candidate TAS identified *in silico* by TASmania database search

The database TASmania [20], which is a discovery-oriented database, was used here to identify candidate TAS in *Mmc* GM12. Our *in silico* search revealed 38 genes encoding candidate toxins or antitoxins (**Table S1**). With respect to our cut-off settings, the following five genes/gene pairs qualified as candidate TAS: MMCAP1_0160/0161, MMCAP1_0890, MMCAP1_0731, MMCAP1_0525, MMCAP1_0753 (**Table 1**). Four of the latter TAS candidates were without a proposed counterpart, the corresponding toxins or antitoxins were manually chosen using ‘guilt by association’, considering the cognate loci of the respective gene as the partner of a two gene operon. A transposon library of JCVI-syn2.0, a synthetic mycoplasma cell, controlled by a reduced genome of *Mmc* strain GM12, suggested JCVISYN2_0132 (referred here as A_132_) and JCVISYN2_0133 (referred to as T_133_) to act as putative TAS [15]. Therefore, we included this candidate TAS into our study.

**Table 1:** Candidate toxins and antitoxins of *M. mycoides* subsp. *capri* GM12 investigated in this study. The top five candidate TAS were identified *in silico* by using the TASmania database.

| Name | Mnemonic/<br>designation in<br>manuscript | Size<br>[nt/aa] | Gene annotation | Pfam/<br>predicted<br>TAS type | E-value in<br>TASmania |
| --- | --- | --- | --- | --- | --- |
| TAS<sub>160/1</sub> | MMCAP2_0160<br>/ T<sub>160</sub> | 753<br>/250 | Abortive<br>infection protein | AbiEii / IV | 1.2e-53 |
|  | MMCAP2_0161<br>/ A<sub>161</sub> | 597<br>/198 | Abortive<br>infection protein | AbiEi / IV | 3.8e-49 |
| TAS<sub>889/0</sub> | MMCAP2_0889<br>/ T<sub>889</sub> | 835<br>/278 | Uncharacterized<br>protein | AbiEii / IV | Guilty by<br>association |
|  | MMCAP2_0890<br>/ A<sub>890</sub> | 600<br>/199 | Conserved<br>hypothetical<br>protein | AbiEi / IV | 2.6e-45 |
| TAS<sub>731/0</sub> | MMCAP2_0731<br>/ T<sub>731</sub> | 1014<br>/337 | Conserved<br>hypothetical<br>protein | AbiEii / IV | 3.8e-29 |
|  | MMCAP2_0730<br>/ A<sub>730</sub> | 600<br>/199 | Conserved<br>hypothetical<br>protein | AbiEi / IV | Guilty by<br>association |
| TAS<sub>525/4</sub> | MMCAP2_0525<br>/ T<sub>525</sub> | 402<br>/133 | Hypothetical<br>protein | MraZ / II | 7e-26 |
|  | MMCAP2_0524<br>/ A<sub>524</sub> | 927<br>/308 | S-adenosyl-<br>methyltransferase | MraW / II | Guilty by<br>association |
| TAS<sub>752/3</sub> | MMCAP2_0752<br>/ T<sub>752</sub> | 573<br>/190 | Cell filamentation<br>protein | Fic / II | Guilty by<br>association |
|  | MMCAP2_0753<br>/ A<sub>753</sub> | 273 /90 | DNA-damage-inducible protein<br>J | RelB / II | 3.5e-9 |
| TAS<sub>133/2</sub> | MMCAP2_0133<br>/ T<sub>133</sub> | 2274<br>/757 | Subtilisin-like<br>serine protease | - / II | N/A |
|  | MMCAP2_0132<br>/ A<sub>132</sub> | 1062<br>/353 | AAA-ATPase | - / II | N/A |

TAS: toxin-antitoxin system, T: toxin, A: antitoxin

The six candidate TAS were distributed throughout the *Mmc* GM12 genome (**Figure 1**). Inspecting the gene loci, we observed that TAS_133/2_ and TAS_731/0_ were arranged in a classical TAS operon structure, with the antitoxin upstream of the toxin and one promotor. TAS_525/4_, TAS_160/1_ and TAS_889/0_ were arranged in a classical operon structure but in antisense orientation. TAS_752/3_ had a non-classical TAS operon structure with the toxin upstream of the antitoxin and an intergenic region of 207 bp. Analysis of this intergenic region using BPROM identified one possible promotor with the putative −10 and −35 boxes, indicating that the antitoxin has its own promotor (**Figure 1**).

**Figure 1:**
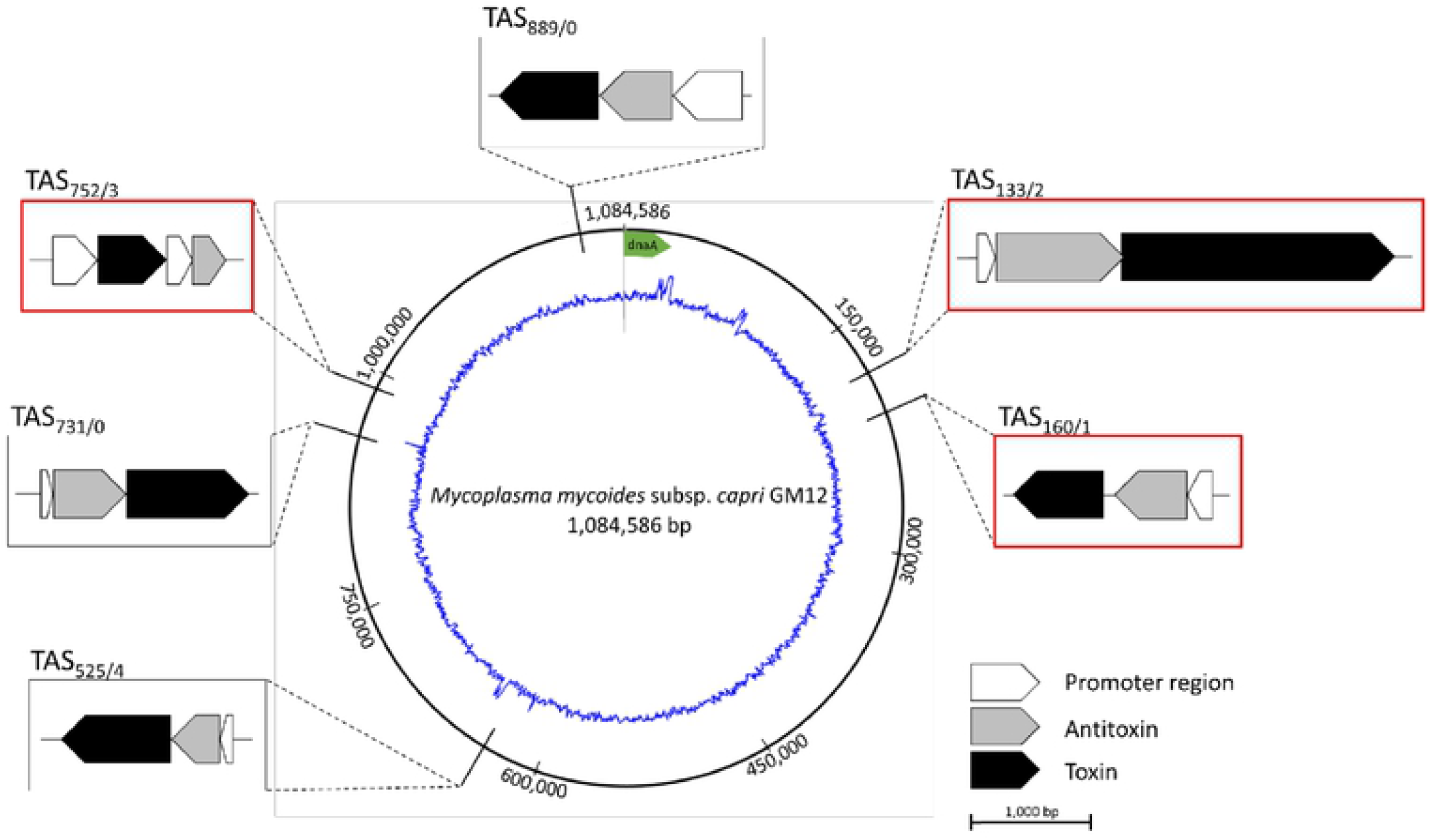
Cartoon displaying the genomic localization and operon structure of candidate TAS of *M. mycoides* subsp. *capri* GM12. The circle displays the genome of GM12 (GenBank accession number: NZ_CP001668.1) including its GC content. Six candidate TAS operons are shown in boxes in relation to their localization in the genome. The promotor region, antitoxin and toxin are marked white, grey and black, respectively. Candidate TAS selected for functionality testing are highlighted with a red box.

Gene description on TASmania database denominated TAS_160/1_ as abortive infection proteins, AbiGII (T_160_) and AbiGI (A_161_), which belong to a well-described protein family (Abi), described to act as toxin-antitoxin systems in e.g. *Streptococcus agalactiae* [21]. T_752_ was denominated as cell filamentation protein (fic) and A_753_ as DNA-damage-inducible protein J.

A_753_ encoded a protein which shares identity with RelB family of TASs, described, amongst others, in *E. coli* [22]. The genes T_133_ and A_132_ were reported to encode an AAA-ATPase and a subtilisin-like serine protease, respectively. A functional TAS (ietAS) with compelling protein similarity exists in *Agrobacterium tumefaciens* [23]. A_890_ and T_731_ were also denominated as belonging to the Abi protein family, according to TASmania and A_525_ was assigned to belong to MraZ protein family, which is similar to the TAS MazEF of *E. coli* [24, 25]. Different types of TAS have been described based on their mode of action and the six TAS identified in our study represent type II and type IV TAS (**Table 1**).

### Detection of transcripts and proteins encoded by candidate TAS

We tested the presence of transcripts covering the genes encoding the six candidate TAS. Therefore, reverse transcription PCR was performed on *Mmc* GM12 RNA with primers targeting the genes encoding candidate toxins, antitoxins and the whole putative two-gene operons. We isolated 24 − 35 μg total RNA from 20 mL cultures. A PCR with primers targeting 16S rRNA did not reveal any amplicons confirming absence of DNA in the RNA preparation. Transcripts of all 6 TAS investigated except for A_890_ were detected via reverse transcription PCR (**Figure S1**).

Afterwards we investigated the presence of candidate toxins and antitoxins using a shot gun proteomics approach, in which we harvested GM12 at one time point, specifically during the log growth phase. Mass spectrometry showed that in *Mmc* GM12 protein abundance of A_524_ is medium, whereas A_132_, A_753_ and A_161_ as well as T_133_ and T_731_ are available in low abundance. We arbitrarily assigned relative abundance thresholds based on the quartile of the log_10_ (mean dNSAF) values distribution: value −3.8 of the lower quartile (Q1) defined as low/medium boundary and value −2.8 of the upper quartile (Q3) defined as medium/high boundary. For reference, house-keeping enzymes dihydrolipoamide S-acetyltransferase, adenylate kinase, glucose-6-phosphate isomerase, DNA-directed RNA polymerase beta chain, guanylate kinase, DNA gyrase subunit B and recombination protein present log_10_ (mean dNSAF) values of −2.0, −2.5, −2.5, −2.6, −2.8, −3.0 and −4.4, respectively. The other candidate toxins and antitoxins were below the detection limit of the mass spectrometry analysis (**Figure 2, Table S2**). In summary only the candidate toxin and antitoxin of TAS_133/2_ have been detected using the shot gun approach, for the TAS.

**Figure 2:**
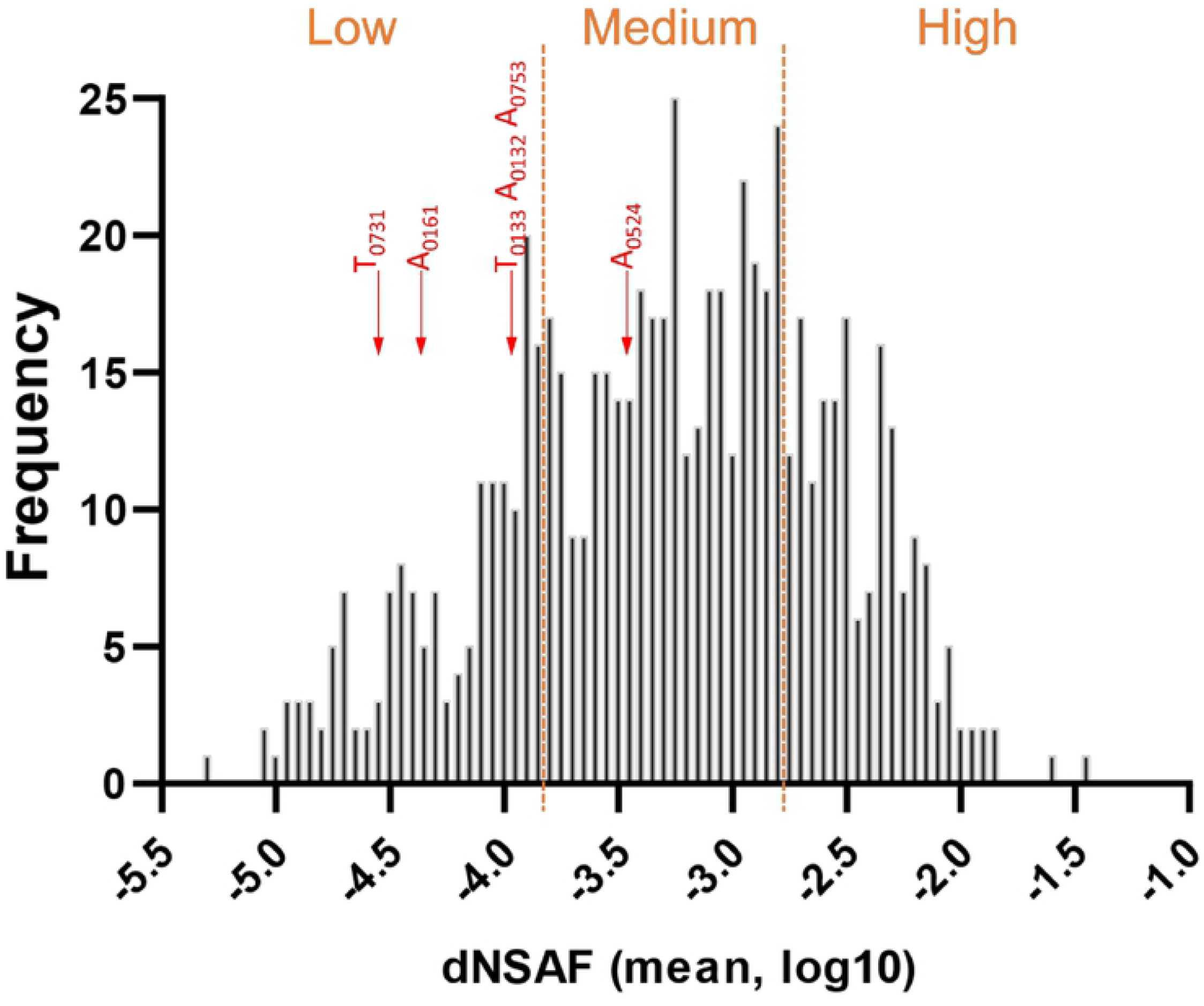
Detection of proteomic signatures of TAS in *in vitro* grown GM12. The histogram shows relative protein abundance in strain GM12 [frequency over distributed normalized spectral abundance factor (dNSAF)], with candidate toxins (T) and antitoxins (A) marked as red arrows and the respective number of the toxin/antitoxin. Toxins and antitoxins had the following dNSAF values: A_524_ −3.47, T_133_ −3.92, A_132_ −3.92, A_753_ −3.92, A_161_ −4.45, T_731_ −4.52. Threshold values of “low, medium and high” expression levels are based on data distribution. Candidate toxins and antitoxins not displayed were below detection limits or absent.

### High conservation in the organization of the *Mmc* genomes

Each of the 15 genomes was assembled into one circularized chromosome and the chromosomes were rotated based on the gene *dnaA*. Key features of each genome are displayed in **Table S3**. The size of the 15 genomes ranged from 1,019,884 to 1,172,410 bp and a G+C content of 23.7 %. All genomes contained 908 (± 93) CDS, 30 tRNAs, two rRNA operons, and 1 tmRNA each. The functional annotation, based on BLASTP of the CDS against a UniProtKB snapshot, revealed an average of 392 hypothetical proteins (∼45% of the total annotated CDS) for each genome. In terms of genome organization, we observed a high level of syntheny **(Figure S2)**. Only strains 152/93 and C260/4 had a large inversion of more than 100.000 bp, which was confirmed by PCRs. Strain 152/93 contained a plasmid with a size of 1.875 bp, which was highly similar to pKMK1 [26] and p*Mmc*-95010 [27] detected in strains GM12 and 95010, respectively.

### Distribution of the six candidate TAS in other mycoplasmas

Next, we were interested to investigate the distribution of the six candidate TAS in different strains of *Mmc* and to extend this analysis to members of the so called ‘*M. mycoides* cluster’, which comprises four additional closely-related ruminant pathogens besides *Mmc*. The presence of genes was investigated by TBLASTN employing complete genome sequences.

Different *Mmc* strains showed that the presence of candidate TAS was not clonal, meaning phylogenetically closely related strains had not necessarily the same set of candidate TAS. Specifically, TAS_752/3_ was present in most strains, except C260/4 and Wi8079, which lacked A_753_. TAS_133/2_ and TAS_160/1_ were only present in strains 7730, My-I, My-325, My-5 and My-18 (**Figure 3A, Table S4**). We revealed that our candidate TAS were also not present in all members of the ‘*M. mycoides* cluster’ (**Figure 3B, Table S4**).

**Figure 3:**
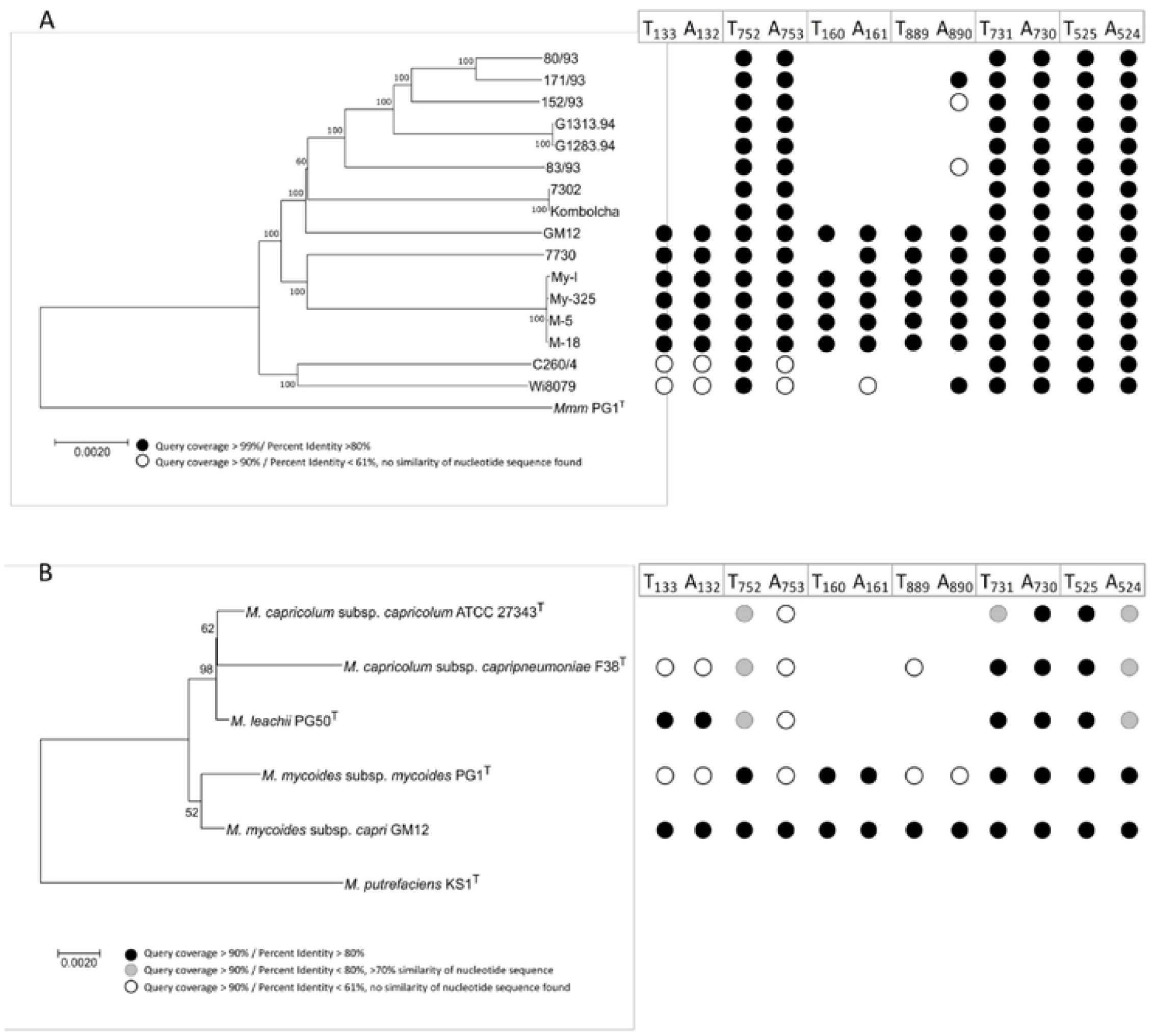
Presence of six GM12 candidate TAS identified *in silico* in other mycoplasmas. Dots on the right indicate the presence of genes encoding candidate toxins or antitoxins. A) Phylogenetic tree based on whole genome sequences of different *M. mycoides* subsp. *capri* (*Mmc*) strains, *M. mycoides* subsp. *mycoides* (*Mmm*) strain PG1^T^ was used as outgroup. The tree was constructed using Neighbor-joining inference method, evolutionary distance was calculated using Maximum Composite Likelihood method with rate variation among site modelled with a gamma distribution. Bootstrap values (100 replicates) are displayed next to tree branches. B) Phylogenetic tree of different members of the ‘*M. mycoides* cluster’ based on 16S rRNA sequences, *M. putrefaciens* strain KS1^T^ was used as outgroup. The tree was constructed using Neighbor-joining inference method. Bootstrap values (100 replicates) are displayed next to tree branches.

### Functionality testing of candidate TAS using heterologous expression

Three of the six candidate TAS were chosen for further characterization. These ones included two candidate TAS identified via TASmania, namely TAS_160/1_ and TAS_752/3_, whereby TAS_160/1_ represented the candidate system with the overall highest score (e-value of 1.2e-53 and 3.8e-49) and TAS_752/3_ represented the candidate system with the lowest score among the five pairs (e-value of 3.5e-9) identified. Moreover, we included the candidate TAS_133/2_ in our studies, since it was suspected to be a TAS because of polar effects of transposon insertions [15]. We aimed to test toxicity of candidate toxins using heterologous expression. Since mycoplasmas lack a cell wall and represent minimal organisms, we opted for testing heterologous expression in different recipients such as *E. coli, B. subtilis* and *M. capricolum*. Consequently we started with *E. coli*, which has been used to confirm TAS in both GRAM-negative and GRAM-positive bacteria [28]. In this bacterial species a bigger toolbox was available than for *Mycoplasma*, like expression vectors with inducible promotors and fast and efficient transformation protocols. Subsequently, genes encoding candidate toxins and antitoxins were cloned into *E. coli* using the established and robust pBAD/His expression system. Heterologous expression confirmed only a negative effect on bacterial growth for the toxin T_133_, while the other two candidate toxins and all three antitoxins tested did not affect the growth of *E. coli* after induction. As expected, growth was only affected when expression of T_133_ was induced, but not when it was repressed. Immunoblotting verified expression of recombinant proteins after induction for all cloned genes (**Figure S3A**). Specifically, four hours post induction, the OD_600_ of *E. coli* transformed with pBAD7His-T_133_ decreased drastically, compared to the various controls (**Figure 4A and Figure S3B**). Interestingly, we observed a phenotype characterized of clumping of *E. coli* cells later in the growth phase in induced *E. coli* cells harboring the vector with T_133_. Then we visualized *E. coli* morphology of those cells using scanning electron microscopy. Visualization revealed that the cell shape started to alter four hours post induction, specifically a fraction of the induced cells was elongated, others were Z-shaped. After seven hours post induction, many cells seem to have burst with only the cell wall remaining visible and a large fraction of cells displayed blebbing at their poles as well as elongation (**Figure 4B**). Empty cell walls clumped together, leading to the clumping observed in liquid cultures by the naked eye. In contrast, cell morphology of all other constructs remained normal (**Figure S3C**). We tested the candidate TAS subsequently in another model organism, namely *B. subtilis*. The rationale behind testing heterologous expression in a GRAM-positive model bacterium was rooted in the closer phylogenetic distance of *B. subtilis* to mycoplasmas that are closely related to the GRAM-positive bacteria and specifically to Clostridia. For these heterologous expression experiments we used the commercial expression system based on pHT01 vector cloned into *B. subtilis* strain 168 Marburg. Construction of the different shuttle plasmids in *E. coli* resulted in six plasmids, each containing a gene encoding either a candidate toxin or antitoxin belonging to the three candidate TAS to be functionally tested. All plasmids except the one containing T_133_ revealed transformants in *B. subtilis*. Since all other plasmids were transformable, toxicity associated with T_133_ is likely to have resulted in absence of viable transformants. Even without IPTG-induction, the promotor of pHT01 causes a low background expression level, which might account for toxic effects of T_133_. Out of the two toxins transformed successfully and tested subsequently in a growth assay, only the toxin T_752_ affected bacterial growth after induction with IPTG, while the other toxin and antitoxins tested had growth curves comparable to *B. subtilis* carrying the empty vector pHT01. Specifically, two hours after IPTG induction, cell density of *B. subtilis* harboring T_752_ started to drop, compared to the other induced toxins and antitoxins, indicating lysis (**Figure 5A, Figure S4A**). Laser scanning micrographs of *B. subtilis* harboring T_752_, showed an altered cell morphology compared to non-induced cells or cells harboring the empty vector (induced or non-induced) (**Figure 5B**), (**Figure S4B**). A large fraction of cells harboring T_752_ burst, shrunk or had holes on the cell surface. Many cells observed during binary fission had one half burst and the other half intact.

**Figure 4:**
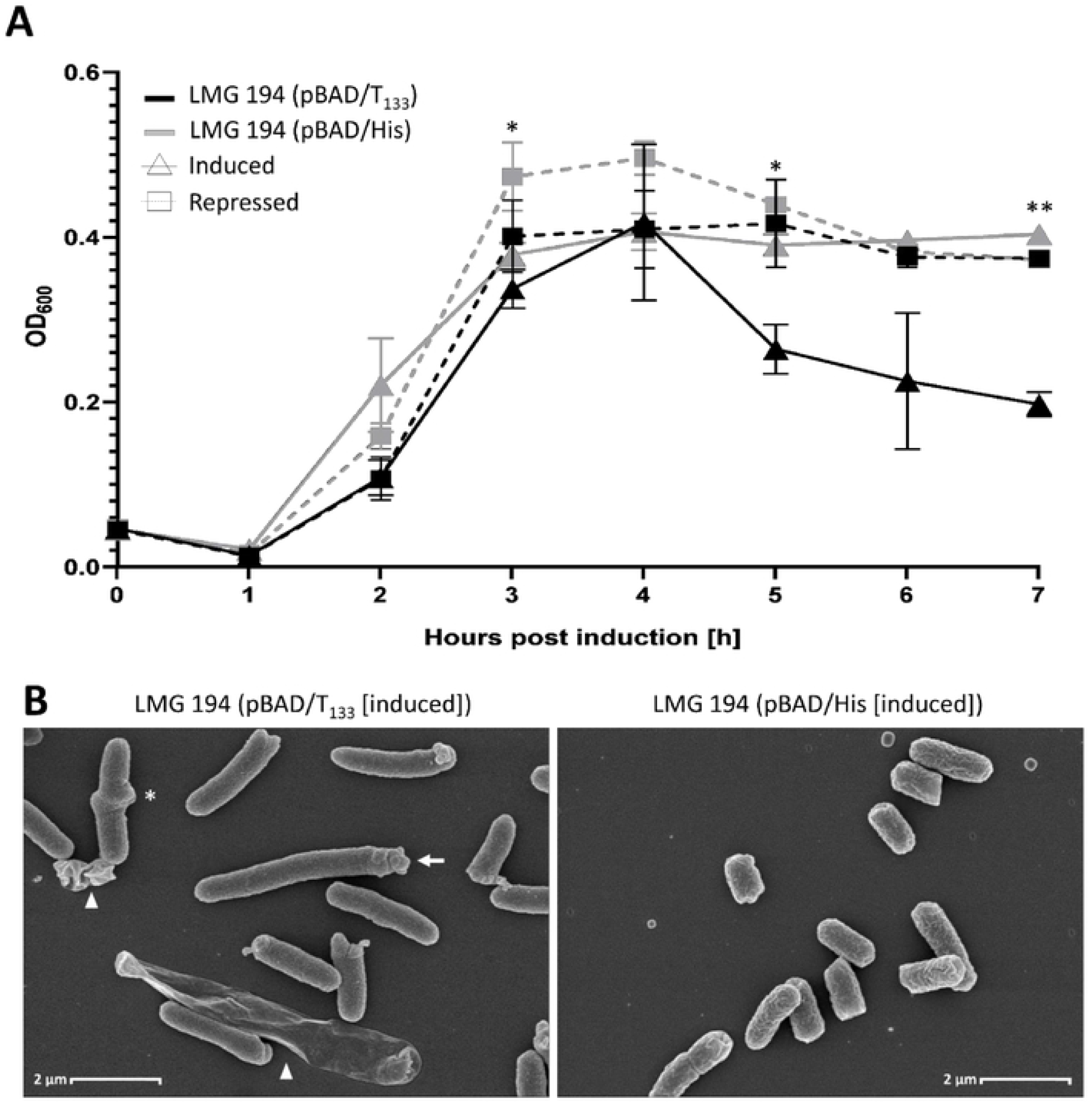
Toxicity of recombinant T_133_ on *E. coli* LMG194. A) Growth curves of *E. coli* LMG194 in response to induction or repression of heterologous expression of T_133_ and empty vector pBAD/His. Each data point represents the mean of three biological replicates, bars indicate standard deviation. The p-values are displayed (* *p* ≤ 0.05, ** *p* ≤ 0.01). B) Scanning electron micrograph (magnification 10,000x) displaying morphological changes of *E. coli* LMG194 after heterologous expression of T_133_. Blebbing at the pole is indicated by an arrow, z-shaped cells by asterisk and cell debris by arrowheads.

**Figure 5:**
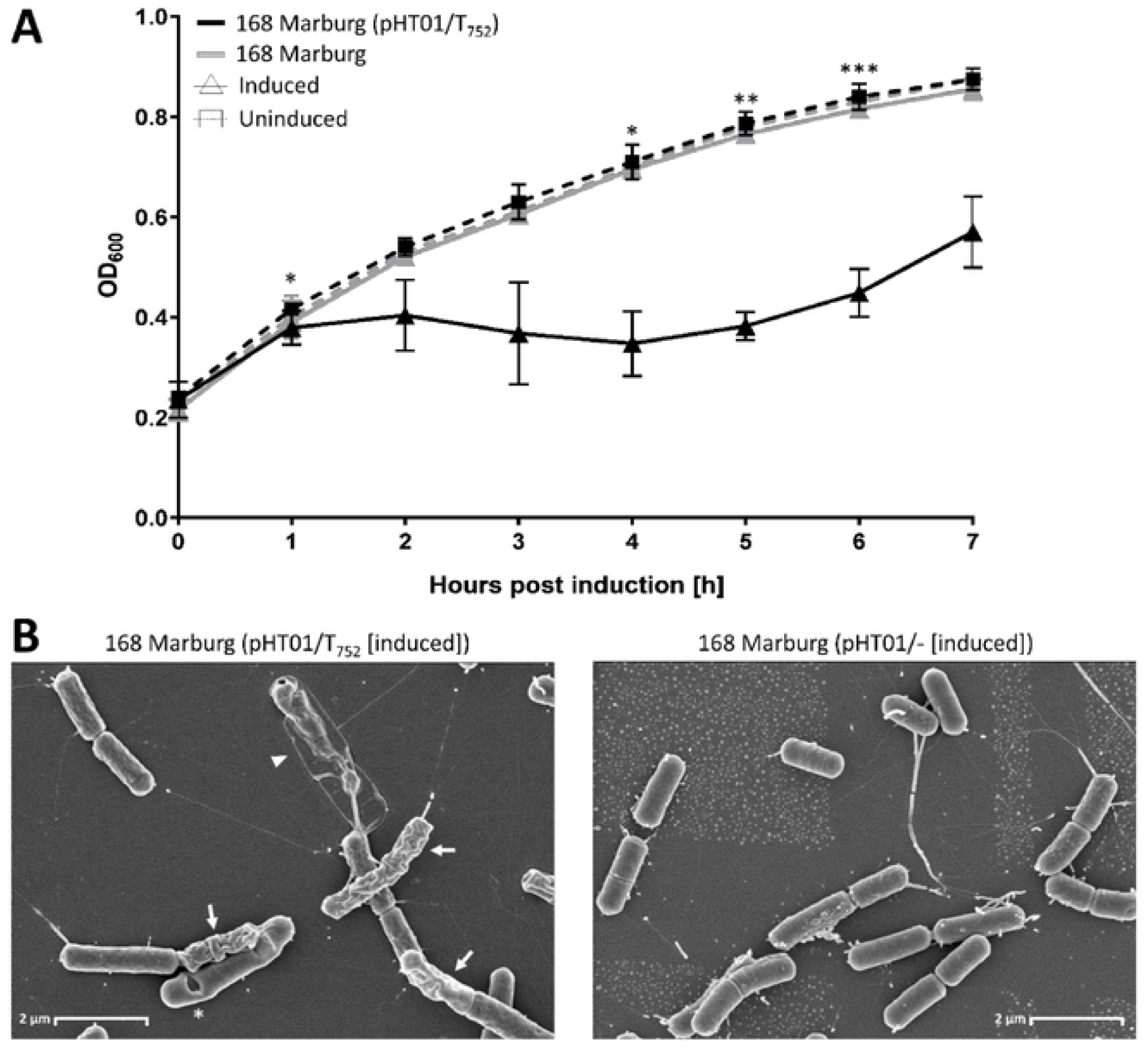
Toxicity of recombinant T_752_ on *B. subtilis* 168 Marburg. A) Growth curves of *B. subtilis* 168 Marburg in response to induction of heterologous expression of T_752_ andstrain 168 Marburg. Each data point represents the mean of three biological replicates, bars indicate standard deviation. The p-values are displayed (* *p* ≤ 0.05, ** *p* ≤ 0.01 and *** *p* ≤ 0.001). B) Scanning electron micrograph (magnification 10,000x) displaying morphological changes of *B. subtilis* 168 Marburg after induction of heterologous expression. Indentations of cells are indicated by asterisk, empty/ shrunk cells by arrowhead and shrunk cells upon division by arrows.

Finally, we took advantage of the availability of stably replicating plasmids available for selected members of the ‘*M. mycoides* cluster’. Since our genome comparisons showed that *Mcap* did not harbor any of the three candidate TAS to be investigated for functionality, *Mcap* represented a “natural knock-out” and was an ideal candidate for heterologous expression. We constructed a set of 4 plasmids for each TAS including the entire TAS under the control of the natural promotor, the antitoxin under the control of the natural promotor and the toxin under the control of the natural promotor as well as the strong spiralin promotor. The cloning of the different plasmids was done via Gibson assembly in *E. coli*. Sequence-verified recombinant plasmids (**Supplementary file S1**) were transformed into *Mcap* and transformation efficacies were monitored to assess toxicity. Transformation of only the antitoxins yielded same amount of transformants as transformation of the empty vector (between 10^5 and 10^6 transformants per µg plasmid). When only the toxins were transformed, 2 logs less transformants were observed compared to the empty vector, which was a significant result. Number of transformants harboring the toxin with the natural promotor compared to the ones containing the spiralin promotor did not differ significantly, except for T_133_ under the control of the spiralin promotor, which did not result in any transformants at all. Transformation rates of the entire TAS did not differ to the empty vector or the individual antitoxins indicating neutralization of the toxic effects by the antitoxin (**Figure 6**).

**Figure 6:**
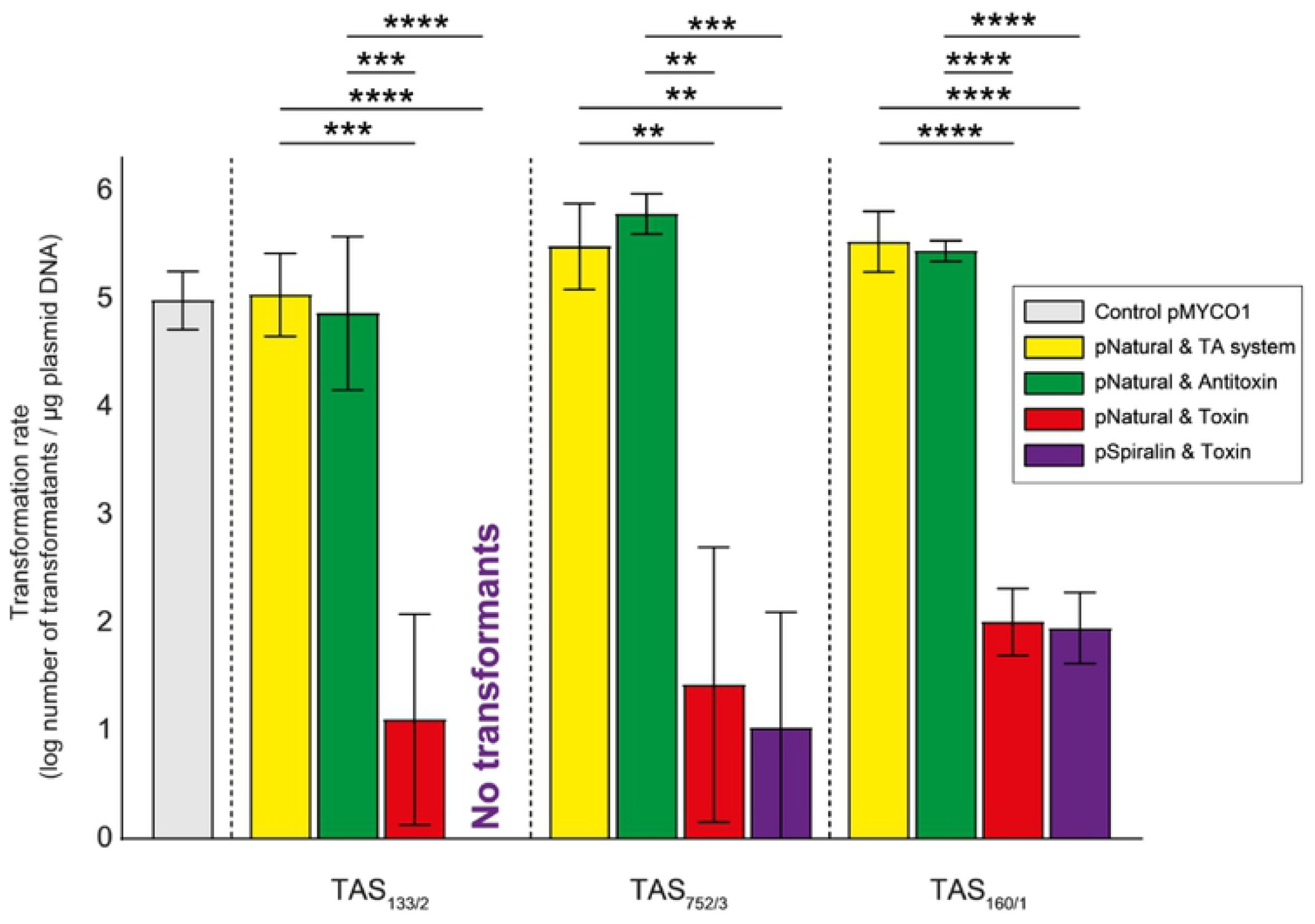
Effect of cloned toxins, antitoxins and TAS on the transformation rate into *M. capricolum* subsp. *capricolum*. *M. capricolum* subsp. *capricolum* ATCC 27343^T^ was transformed with different plasmid constructs harboring either entire TAS operons (TAS_133/2_, TAS_752/3_ and TAS_160/1_), individual toxins or antitoxins. Empty pMYCO1 plasmid was used as a positive control. Each column represents the mean of three independent biological replicates and bars indicate standard deviations. Significance is indicated (** *p* ≤ 0.01, *** *p* ≤ 0.001, **** *p* ≤ 0.0001).

Assessment of vector and insert presence was tested in 3-5 transformants per transformation experiment. It confirmed the presence of antitoxins and entire TAS as expected. *Mcap* cells transformed with the toxin T_133_ under the control of the natural promotor, were found to contain at most partial toxin sequences. We confirmed the presence of T_752_ sequence with its natural promotor in a fraction of the transformants investigated, but transformants of the toxin under the control with the spiralin promotor did not contain T_752_ sequences. The sequence of T_160_ were found in all the examined clones with natural and spiralin promotor.

### Donor-recipient phylogenetic distance impacts functional confirmation of TAS

Toxicity testing has been carried out in the three recipient species *E. coli, B. subtilis* and *Mcap* listed from most to least phylogenetic distance to the TAS donor species *Mmc*. We started testing the two components of the TAS individually in *E. coli* using the established expression system based on pBAD/His. Only T_133_ conferred toxic activity to its recipient upon induction of expression, indicating TAS_133/2_ to be functional. Next, we proceeded to test the systems in *B. subtilis*, which is phylogenetically much closer to *Mmc* than *E. coli*, but likewise consists of a much bigger proteome and a cell wall, factors that are likely to interfere with TAS functionality. The pHT01 vector used has some basal expression, since its promotor is not tight. It was not possible to transform the vector containing T_133_ into *B. subtilis*, indicating toxicity affecting transformation. Additionally, the growth of the clone carrying T_752_ was impacted after induction of expression. Therefore, two out of three TAS tested functional in *B. subtilis*. In *Mcap* all TAS tested functional and moreover the antitoxins neutralized the toxins. In summary, heterologous expression systems confirmed functionality influenced by donor-recipient phylogenetic distance.

## Discussion

This study aimed to identify and to produce proof of the functionality of toxin antitoxin systems (TAS) in minimalistic mycoplasmas for the first time. Therefore, we employed our model organism, the highly pathogenic *Mycoplasma mycoides* subsp. *capri* strain GM12. Many pathogenic mycoplasmas are associated with chronic infections, which theoretically suggests a role for TAS to support existence in a stealth mode that protects the mycoplasmas from immune responses. In fact, live *Mycoplasma mycoides* subsp. *mycoides*, the causative agent of contagious bovine pleuropneumonia, have been isolated from lung sequestra >6 months after experimental infection [10] and the same mycoplasmas survived even in the presence of high antibody titers against the latter. The *in silico* analysis of strain GM12 revealed 18 chromosomally encoded putative candidate TAS elements of which five candidate TAS with the lowest TASmania output e-values were short-listed for our study. In *Mycobacterium tuberculosis*, a deathly human respiratory pathogen with a much larger genome, at least 79 TAS have been reported to be involved in numerous mechanisms regarding cellular processes such as cell death, persistence and virulence [29-31]. In contrast, *Campylobacter jejuni*, a human pathogen causing diarrhea, has been reported to contain no more than three TAS, which underlines a great variety in the abundance of TAS in different prokaryotic genera [29]. So far, functional TAS in mycoplasmas have never been convincingly shown but only indicated *in silico* based on sequence similarity to other TAS [32]. In this study, we focused on the characterization of the TAS with the lowest e-values, since they provided the biggest odds of having a functional TAS. We added the candidate TAS_133/2_, which was proposed recently [15] based on the analysis of a transposon insertion library of JCVI-syn2.0, a synthetic cell controlled by a drastically reduced genome [2]. These six candidate TAS were grouped into types II and IV (**Table 1**) systems. In type II systems the proteinaceous antitoxin is binding to the toxin to prevent its action, in type IV systems the antitoxin competitively binds the toxin cell target. First, we aimed to identify transcripts and protein signatures of the six systems during in vitro growth indicating a biological function of the genes investigated. We identified transcripts of all TAS elements except for A_890_ supporting a role of transcribed genes in the physiology of GM12. The absence of a transcript for A_890_ does not exclude its transcription, since low levels of transcription and a high turnaround time of transcripts can cause transcripts to be below the detection limit. Protein signatures encoded by the six TAS elements were detected for the entire TAS_133/2_ and the individual elements of four TAS, namely A_161_, A_524_, T_731_ and A_753_. The only system we had no proteomic signatures was the TAS_889/0_ for which we did not detect a transcript of A_890_. The detection of proteomic signatures in five out of six candidate TAS using a generic proteomics approach indicates a high probability of functionality especially for the TAS_133/2_. The complete genomes of 16 *Mmc* strains revealed a high level of synteny, only two strains had a large inversion in contrast to the other strains. Interestingly, three out of six TAS were present in all 16 *Mmc* genomes investigated, while phylogenetic clades sometimes also had different TAS fingerprints (**Figure 3A**) making horizontal gene transfer likely to have acted on TAS acquisition or loss. Investigation of genomes of the five members of the *M. mycoides* cluster, which consists of three species (two species contain two subspecies) [33], also confirmed the presence of TAS_752/3_, TAS_731/0_ and TAS_525/4_ in all species investigated (**Figure 3B**) pointing towards old rather conserved acquisitions and a certain plasticity of the genome with respect to TAS. The ‘*M. mycoides* cluster’ is evolutionary young, since its common ancestor dates back 10,000 years [33] coinciding with the start of ruminants’ domestication. Therefore, these different TAS fingerprints likely formed within the last 10,000 years and horizontal gene transfer [34-36] is likely to have contributed to it.

Next, we tested functionality and shortlisted three candidate TAS for this purpose, based on at least one of the following arbitrary criteria i) a low e-value (<10e-7), ii) TAS-related name for the nearest Pfam annotation of the TASmania HMM profile that hit the locus, iii) TAS-related gene description and iv) presence of candidate toxin and cognate antitoxin in our proteomic analysis. We selected TAS_133/2_, since protein signatures were reported for its toxin and cognate antitoxin, TAS_160/1_, since it was the type IV system with the highest TASmania score and TAS_752/3_, and since it was suggested to be a type II system and its toxin and cognate antitoxin encoding gene had gene annotations. For functional testing we established a model organism-based approach by expressing recombinant toxins and antitoxins in the different recipients *E. coli* and *B. subtilis* employing expression plasmids with inducible promotors as well as transforming the candidates in the closely related *Mycoplasma capricolum* subsp. *capricolum*. We started to test toxicity of candidate toxins in *E. coli*, which has been done for other candidate TAS of GRAM-positive bacteria, e.g. Streptomyces [37] or Mycobacteria [28]. Unexpectedly, although being high score candidates, two out of three candidate TAS did not test positive in *E. coli*. Only recombinant T_133_, annotated as subtilisin-like serine protease, had a toxic effect on the growth of *E. coli* LMG194. Serine proteases cleave peptides or proteins. A role of serine proteases as toxins in TAS has not been reported except for IetS [23] in the GRAM-negative plant pathogen *Agrobacterium tumefaciens*, a toxin which was shown to be involved in plasmid stability. Heterologous expression of T_133_ in *E. coli* resulted in cell clumping in liquid culture and elongated and filamented cells. The observed morphological changes can be attributed to either direct toxic effects or a stress response of *E. coli* towards the recombinant T_133_, which should be investigated in future studies. Cell filamentation has been reported to occur as an SOS response upon DNA damage in other bacteria [38, 39]. Heterologous expression in *B. subtilis* revealed toxicity of recombinant T_133_ (plasmid carrying T_133_ was not transformable) and of T_752_, which had a toxic effect on the growth. The latter encodes for a fic protein, which are known in bacteria to disrupt the DNA topology via adenylation and causes growth arrest [40]. Finally, functionality of all three TAS investigated was shown in the transformation experiments carried out with *Mcap*. These experiments clearly also showed the capacity of the antitoxins to neutralize the effects of the toxins. Interestingly, the type IV system TAS_160/1_ only showed functionality in *Mcap*, probably due to the fact that in such systems toxin and antitoxin bind to the same target, which was probably only available in a mycoplasma.

In conclusion, this study identified six candidate TAS and tested functionality of three out of the six chromosomally encoded TAS in *Mmc* GM12. Functionality of the three candidate TAS was shown, as expression of only the toxin results in perturbations of cell vitality or in cell death whereas the cognate antitoxin compensated the toxin-induced effects completely. Further, we evaluated different model organisms as *E. coli, B. subtilis* and *Mycoplasma capricolum* for heterologous expression of TAS. According to our results, toxin action seems to be dependent on phylogenetic proximity towards the bacterial species from which candidate TAS originate. TAS patterns were different in different *Mycoplasma* species and subspecies indicating a level of plasticity of the mycoplasmas’ genomes with respect to TAS. Since mycoplasmas are minimal organisms [11] amendable to synthetic genomics techniques [41], their host-pathogen interactions can be investigated in the native host [17, 18, 42] and as they contain only a limited number of TAS, they represent an ideal model organism to study the role of TAS in pathogenicity.

## Materials and methods

### *In silico* identification of candidate TAS

Identification of candidate TAS in *Mmc* strain GM12 [16] was performed by employing the TASmania discovery pipeline [20], which uses data mining on the Ensembl Bacteria database [43] to identify candidate toxin and antitoxin proteins. The e-value was set to the default value of 1e-05. TAS candidates to be characterized in the study had the following minimal requirements: an e-value lower than 1e-07, a protein family (pfam) describing known TAS and gene description indicating possible candidates. If TAS candidates were without a counterpart (toxin or antitoxin), the counterparts were manually selected using ‘guilt by association’, considering the cognate loci of the respective gene as the partner of a two gene operon. Intergenic regions close to toxin and antitoxin genes on the *Mmc* GM12 genome were analyzed using BPROM (Softberry) [44] to identify putative promotor regions.

### Bacterial strains and plasmids used for cloning in this study

Plasmid based on pBAD/His (Invitrogen) or pHT01 (MoBiTech) and pMYCO1 were constructed in *E. coli* DH5α (Takara) and Stellar™ (Takara), respectively. Heterologous expression based on pBAD/His and pHT01 was done in *E. coli* LMG194 (Invitrogen) and *B. subtilis* 168 Marburg (MoBiTech), respectively. The restriction-free *M. capricolum* subsp. *capricolum* mutant strain *M. capricolum* RE(–) [41] was used as recipient for transformation experiments involving pMYCO1-derived plasmids [45].

### Detection of transcripts for candidate TAS

*Mycoplasma mycoides* subsp. *capri* (*Mmc*) strain GM12 was grown in 10 ml SP5 medium at 37°C until the culture reached pH 6.5. Subsequently, the culture was centrifuged at 4,255 x *g* at 10°C for 15 min. The supernatant was discarded and the bacterial cell pellet was used for RNA isolation. RNA was isolated using the Zymo Research Quick-RNA Fungal/Bacterial Miniprep™ kit. Briefly, 800 μl of RNA Lysis Buffer were added to the pellet and the protocol was followed starting at step 4 of the manual. After isolation, a DNase treatment was performed using the Clean & Concentrator™ −5 kit (ZymoResearch), omitting the first step of the protocol (addition of Binding Buffer) and using the IICR columns provided in the kit. Reverse transcription PCR was performed using SuperScript™ IV RT Mix (Invitrogen) and 2x Platinum™ SuperFi™ RT-PCR Master Mix (Invitrogen) using primers No. 59-82 listed in **Table S4**. In brief, 12.5 μl of 2x Platinum™ SuperFi™ RT-PCR Master Mix, 0.25 μl SuperScript™ IV RT Mix, 1.25 μl per primer (10 μM), 1μl DNase treated RNA (∼ 32 ng) and water up to a volume of 25 μl were mixed per sample. Amplicons were visualized on a 1% agarose gel.

### Proteomic analysis

Strain GM12 was grown in 50 mL of SP5 media until mid to late logarithmic phase (pH of media at harvest was 6.5) and centrifuged at 3,000 x *g* for 20 min. Cells were washed three times with PBS and pellets were kept at −80 for subsequent analysis. The protein concentration was measured using the Pierce BCA Protein Assay Kit (ThermoScientific) according to vendor’s instructions. Samples for mass spectrometry analysis were prepared following standard protocols [46]. Briefly, cell pellets were thawed on ice and lysed in 8M urea/100mM Tris-HCl; proteins were precipitated in cold acetone over-night and denatured with 8M urea/50mM Tris-HCl, then digested with trypsine over-night at room-temperature in 1.6M urea /20mM Tris-Hcl /2mM CaCl_2_. Enzymatic digestion was stopped by adding 1/20-volume of 20% (v/v) tri-fluoroacetic (TFA). The digests were analyzed by liquid chromatography (LC)-MS/MS (PROXEON coupled to a QExactive HF mass spectrometer, ThermoFisher Scientific). Samples were further processed against custom databases by Transproteomics pipeline (TPP) tools [47]. Four database search engines were used: Comet [48], Xtandem [49], MSGF [50] and MyriMatch [51]. Each search was followed by the application of the PeptideProphet tool [52]; the iProphet [53] tool was then used to combine the search results, which were filtered at the false discovery rate of 0.01; furthermore, the identification was only accepted if at least two of the search engines agreed on the identification. The decoy approach was used for such custom databases containing standard entries. Protein inference was performed with ProteinProphet. For those protein groups accepted by a false discovery rate filter of 0.01, a Normalized Spectral Abundance Factor (NSAF) [54] was calculated based on the peptide to spectrum match count; shared peptides were accounted for by the method published elsewhere [55].

### Cloning of genes encoding candidate toxins and antitoxins in *Escherichia coli*

*E. coli*-codon optimized synthetic genes encoding candidate toxins or antitoxins were synthesized and cloned into pUC57 and sequence-verified by GenScript (**Supplementary File S1)**. For ligation into the expression vector pBAD/His, genes to be cloned, were PCR-amplified using primers that contained restriction sites for *EcoRI* and *SacI* (**Table S5**, primers No. 1-6) and the high-fidelity polymerase Q5 (New England Biolabs) according to vendor’s manual. Amplicons were purified using the High Pure PCR Product Purification Kit (Roche) and afterwards restricted with *EcoRI* (New England Biolabs) and *SacI* (New England Biolabs) followed by another purification using the above-mentioned kit. Toxin genes to be cloned were retrieved from the plasmids that contained the synthetic genes via restriction followed by gel purification. Restricted fragments to be cloned were ligated into the *EcoRI* and *SacI* restricted pBAD/His using T4 Ligase (Promega) before being heat shock-transformed into chemically competent *E. coli* DH5α (Takara) or Stellar™ (Takara) using standard protocols [56]. Recombinant plasmids from transformants were isolated using the QIAprep® Spin Miniprep Kit (Qiagen) and sequence-verified via Sanger sequencing (Microsynth) with specific primers (**Table S5**, primers No. 7-8). Sequence-verified plasmids were heat shock-transformed into chemically competent *E. coli* LMG194 for subsequent expression. Transformants were verified as described above and stored at −80 °C for subsequent experiments.

### Functional testing of candidate *Mycoplasma* TAS using heterologous expression in *E. coli*

Growth curves of *E. coli* LMG194 containing inducible expression vector pBAD/His carrying genes encoding individual candidate toxins and antitoxins were performed in 96-well plates (Tissue culture test plates, 92096, TPP) and monitored over a period of seven hours. All clones were cultured in LB medium supplemented with 50 μg/mL ampicillin. Twenty µL of overnight cultures of clones to be tested were mixed with 180 μL LB supplemented with either 0.2% (w/v) arabinose or 0.2% (w/v) glucose to induce or repress protein expression, respectively.

Cultures in 96-well plates were incubated at 37 °C under agitation (500 rpm) using a ThermoMixerC (Eppendorf). Optical density was measured at 600nm using the VERSAmax plate reader (Bucher Biotec, Basel, Switzerland). Final datasets were obtained from three biological replicates (each biological replicate consisted of three technical replicates). To assess the significance of the heterologous expression on growth of the clones at every time point, we applied a three-way (time, treatment, construct) repeated-measures ANOVA and decomposed into post hoc tests based on pairwise comparisons with a Bonferroni adjustment of the p-value. The statistical analysis was done using R version 3.6.2.

### Detection of recombinant proteins using immunoblots

Heterologous expression in *E. coli* was verified by immunoblot using anti-HIS antibodies tagging the recombinant fusion proteins. Briefly, 200 µL of overnight culture were mixed with 20 ml of LB supplemented with 50 µg/mL ampicillin and arabinose (final concentration 0.2%).

Cultures were grown on a shaking incubator at 220 rpm (Lab-Shaker) at 37 °C and samples of 1 mL were sequentially removed at several time points for subsequent analysis. As controls served the same clones grown in the same media except that arabinose was exchanged with glucose (final concentration 0.2%). Samples were spun down at 16,100 x *g* at room temperature for 3 min. The supernatants were discarded, the pellets were resuspended in PBS (Merck) and the protein concentration was determined using the Pierce™ BCA Protein Assay Kit (ThermoScientific). Immunoblotting was basically carried out as described recently [18]. Briefly, 0.1 mg of proteins in the cell pellet were separated onto a 12% SDS-PAGE gel using standard procedures [56]. Size separated proteins were transferred onto a 0.2 μm pore-size nitrocellulose membrane (BIO RAD) using the Trans-Blot® Turbo Transfer System (BIO RAD) at 25 volts, 1.0 A for 30 min. PBS supplemented with 0.1% Tween-20 (Merck) and 2% BSA (Sigma) served as blocking buffer and monoclonal mouse-derived anti-HIS antibody (LS-C57341, LsBio) and HRP conjugated goat anti-mouse IgG antibody (AP308P, Sigma) served as primary and secondary antibodies, respectively. Primary antibodies were diluted in blocking buffer at 1:1,000 and incubated with the membrane for 1 hour, afterwards the membrane was washed 3 times in PBS supplemented with 0.1% Tween-20 for 10 min. Then the membrane was incubated with the secondary antibody diluted at 1:70,000 in blocking buffer for 1 hour. After three washes as done before the membrane was incubated with SuperSignal™ West Pico PLUS Chemiluminescent Substrate (ThermoScientific) according to manufacturer’s recommendations. Chemiluminescence was visualized with the CCD-camera Fusion FX (Vilber) and analyzed by the software Evolution.

### Cloning of genes encoding candidate toxins and antitoxins in *Bacillus subtilis*

Synthetic genes to be cloned into *B. subtilis* were first cloned into *E. coli* using the shuttle vector pHT01 (MoBiTec). Therefore, genes were PCR-amplified using synthetic genes cloned into pUC57 (see above) as template, primers that contained restriction sites for *BamHI* and *XmaI* or *XbaI* (**Table S5**, primers No. 9-16 and 95-98) and the high-fidelity polymerase Q5 (New England Biolabs) according to vendor’s manual. Amplicons were purified using the High Pure PCR Product Purification Kit (Roche) and afterwards restricted with *BamHI* (ThermoScientific) and *XmaI* (ThermoScientific) or *XbaI* (ThermoScientific) followed by another purification step using the above-mentioned kit. The restricted amplicons were ligated into the *BamHI* and *XmaI* (*BamHI* and *XbaI*) restricted pHT01 using T4 Ligase (Promega) before being heat shock-transformed into chemically competent *E. coli* Stellar™ (Takara) following standard protocols [56]. *E. coli* Stellar™ was grown on LB media supplemented with ampicillin at 50 µl/mL. Colony PCR was performed with primers No. 17-24 and 99-102 (**Table S5**). Recombinant plasmids from transformants were isolated using the QIAprep® Spin Miniprep Kit (Qiagen) and sequence verified via Sanger sequencing (Microsynth) with specific primers (**Table S5**, No. 25-26). Sequence-verified plasmids were transformed into *Bacillus subtilis* 168 Marburg (MoBiTec) using vendor’s protocols. Therefore, we followed the protocol for making naturally competent cells. Transformants were selected on agar plates containing 5 µg/mL chloramphenicol. Transformants were lysed as described elsewhere [57] and presence of plasmids was confirmed via PCR targeting cloned genes. Clones were stored at −80 °C for subsequent experiments.

### Functional testing of candidate TAS using heterologous expression in *B. subtilis*

Growth curves of *B. subtilis* 168 Marburg containing inducible expression vector pHT01 carrying genes encoding individual candidate toxins and antitoxins were performed in 96-well plates (Tissue culture test plates, 92096, TPP) and monitored over a period of seven hours. All clones were cultured in LB medium supplemented with 5 μg/mL chloramphenicol (Serva). Overnight cultures of *B. subtilis* 168 Marburg were diluted to an OD_600nm_ of 0.15 in a total volume of 5 mL, which was subsequently incubated at 37 °C. When cultures reached an OD_600_ of 0.7-0.8, IPTG (Roche Diagnostics) was added to a final concentration of 1 mM for induction of heterologous protein expression. Cultures in 96-well plates were incubated at 37 °C under agitation (500 rpm) using a ThermoMixerC (Eppendorf). Optical density was measured at 600 nm using the VERSAmax plate reader (Bucher Biotec). Final datasets were obtained from three biological replicates (each biological replicate consisted of three technical replicates).

To assess the significance of the heterologous expression on growth of the clones at every time point, we applied a three-way (time, treatment, construct) repeated-measures ANOVA and decomposed into post hoc tests based on pairwise comparisons with a Bonferroni adjustment of the *p*-value. The statistical analysis was done using R version 3.6.2.

### Cloning of genes encoding candidate toxins and antitoxins into *Mycoplasma capricolum* subsp. *capricolum*

The three candidate TAS operons, as well as each of the genes encoding candidate toxins and antitoxins, were individually cloned into pMYCO1. Candidate genes or operons were PCR amplified using Q5 high fidelity DNA polymerase (New England Biolabs) according to manufacturer’s instructions using primers (No. 27-49) listed in **Table S5**. DNA assembly was performed in *E. coli* using the NEBuilder HiFi DNA Assembly kit (New England Biolabs) following manufacturers’ instructions. When necessary, the natural promoters of each of the TAS were amplified independently and positioned in frame with their respective candidate toxins. Similarly, the spiralin promoter was amplified from the plasmid pMT85tetM-PSlacZ-pRS313 (GenBank accession number KX011460) and assembled upstream of each toxin. All the constructions were sequenced using primers No. 57 and 94 (**Table S5**).

### Functional testing of candidate TAS using heterologous expression in *M. capricolum* subsp. *capricolum* (RE-)

*M. capricolum* subsp. *capricolum* (*Mcap*) (RE-) was selected to probe functionality of the three candidate TAS as none of the latter appeared to be present in its genome (**Figure 3B**). All 12 pMYCO1-based constructs were transformed into *Mcap* (RE-) as previously described [58] using 1 μg of plasmid DNA. Three biological replicates including three technical replicates per biological replicate were carried out. Transformants were passaged for three consecutive rounds in 1ml SP5 medium supplemented with 5 µg/mL tetracycline and incubated at 37 °C to confirm the resistant phenotype. Transformants were screened using primer pairs specific for the inserted TAS elements (**Table S5**, No. 50-58). Numbers of transformants were used to analyse transformation efficacies. An ordinary one-way ANOVA test with Tukey’s multiple comparison test was applied (GraphPad Prism version 8.0.0, San Diego, California USA).

### Scanning electron microscopy (SEM) of *E. coli* and *B. subtilis* recombinant clones

The morphology of *E. coli* and *B. subtilis* cells upon toxin induction was investigated by scanning electron microscopy. Clones to be visualized were cultured and expression of the candidate toxins was induced as described above. Samples collected during expression (1 ml) were centrifuged at 1,500 x *g* at room temperature for 3 min. The pellets were washed 3 times with 1 ml PBS and finally resuspended in 250 μl PBS. Subsequently, 250 μl of 5% glutaraldehyde (Merck) in 0.2M cacodylate buffer, pH 7.4, was slowly added to the samples with gentle agitation. The tubes were stored at 4 °C until further processing. Fixed cells were cytospun onto platinum-sputtered and PLL-coated coverslips (40 μl at 125 x *g*, 5 min). The coverslips were then washed 3 times with 0.1M cacodylate buffer, pH 7.4, postfixed in 1% OsO4 (Polysciences) in 0.1M cacodylate buffer for 30 min, washed 3 times again and dehydrated in an ascending series of ethanol (70%, 80%, 94%, 100%, 100%, 100% ethanol at room temperature for 15 min each). Samples were dried by evaporation of Hexamethyldisilazane (Merck). Samples were coated with 15 nm of platinum in a high vacuum coating unit (CCU-010; Safematic) and examined with a scanning electron microscope DSM 982 Gemini (Zeiss) at an accelerating voltage of 5 kV at a working distance of 4 mm.

### Isolation of genomic DNA from for next generation sequencing

Mycoplasmas grown in 8 ml of SP5 medium overnight were pelleted at 4,255 x *g* at 10 °C for 15 min. The supernatant was discarded and gDNA was extracted using the Wizard Genomic DNA purification kit (Promega) according to vendor’s protocol. Plasmid DNA was extracted from 10 ml overnight culture using QIAprep® Spin Miniprep Kit (QIAGEN) with the following adaptations: additional washing with buffer PB and using the double of the recommended volume of buffers P1, P2 and P3. DNA concentration and purity were determined by agarose gel separation, Nanodrop (ND-1000 Spectrophotometer, Witec AG) and Qubit Fluorometric quantification (Invitrogen).

### Next generation sequencing of *Mycoplasma mycoides* subsp. *capri* strains

Sequencing was carried out at the Lausanne Genomic Technologies Facility using the PacBio. Briefly, DNA was sheared in a Covaris g-TUBE (Covaris, Woburn, MA, USA) to obtain 10 kbp fragments. After shearing, the DNA size distribution was checked on a Fragment Analyzer (Advanced Analytical Technologies, Ames, IA, USA). A barcoded SMRTbell library was prepared using 480 ng of gDNA using the PacBio SMRTbell Template Express Prep Kit 2.0 (Pacific Biosciences, Menlo Park, CA, USA) according to the manufacturer’s recommendations. Libraries were pooled and sequenced with v3.0/v3.0 chemistry on a PacBio Sequel instrument (Pacific Biosciences, Menlo Park, CA, USA) at 10 hours movie time, pre-extension time of 2 hours, using one SMRT cell v3.

### Genome assembly, annotation and alignments

Genomes were assembled from PacBio reads using the software Flye, version 2.6 [59]. Circularized genomes were polished with three rounds with the software Arrow [single-molecule real-time (SMRT) Link version 8 package]. Plasmids were Sanger sequenced after cloning into *E. coli* using pUC19. Genomes were rotated to the first nucleotide of the start codon of the *dnaA* gene. Sequences were then annotated using Prokka, version 1.13 [60]. The program Mauve (version 20150226) [61] was used for the construction of the genome alignment of multiple *Mycoplasma* strains.

### Generation of a phylogenetic tree of members of the ‘*M. mycoides* cluster’

The phylogenetic tree was generated using 16S rRNA sequences retrieved from GenBank. The following strains were included in the analysis: *Mycoplasma mycoides* subsp. *capri* strain GM12 (GenBank accession number NZ_CP001668.1), *Mycoplasma mycoides* subsp. *mycoides* strain PG1^T^ (GenBank accession number BX293980.2), *Mycoplasma capricolum* subsp. *capricolum* strain California kid^T^ (GenBank accession number NC_007633.1), *Mycoplasma capricolum* subsp. *capripneumoniae* strain F38^T^ (GenBank accession number LN515398.1), *Mycoplasma leachii* strain PG50^T^ (GenBank accession number NC_014751.1) and *Mycoplasma putrefaciens* strain KS1^T^ (GenBank accession number NC_015946.1). The software MEGA, version 10.0.5 [62] was used for construction of phylogenetic trees. For the phylogenetic tree displaying the *M. mycoides* cluster, evolutionary history was inferred using the Neighbor-Joining method and evolutionary distance was computed with Kimura 2-parameter method. For statistical support of the phylogeny, 100 replicates were used for the bootstrap test.

### Generation of phylogenetic trees of *Mmc* strains using whole genome sequences

For tree generation, whole genome sequences of *Mycoplasma mycoides* subsp. *capri* strains were used and *Mycoplasma mycoides* subsp. *mycoides* PG1^T^ was included as an outgroup. Mapping of genomes was performed by the Reference sequence Alignment based Phylogeny builder (REALPHY) [63]. For tree construction, the software MEGA (version 10.0.5) [62] was used. Evolutionary history was inferred using the Neighbor-joining method. Evolutionary distances were computed with Maximum Composite Likelihood method. Rate variation among sites was modelled with a gamma distribution.

### Graphic representation of toxin and antitoxin presence on genomes of mycoplasmas

A graphic representation of gene presence was generated based on TBLASTN similarity of candidate toxin and antitoxin amino acid sequences and genomes of different mycoplasmas. The presence and conservation of each toxin and antitoxin candidate was assessed using TBLASTN [64] with *Mmc* GM12 amino acid sequences of candidate toxins and antitoxins as query against whole genome sequences of different *Mycoplasmas* and *Mmc* strains. All sequences with query coverage >99% and percent identity >80% were considered as being present on the respective genome. If Query coverage was <55%, we did assume the candidate toxin/antitoxin does not exist in the respective genome. In case the query coverage was >70% but percent identity was <61%, a BLASTN was additionally performed to check for nucleotide sequence similarity. If the nucleotide sequence identity query coverage of BLASTN was >80% and percent identity >70%, we assumed the candidate toxin/antitoxin to be present on the respective genome.

## Author contributions

JJ designed the research. VH, FL, RSB, HAE, MHS performed the research. VH, FL, HAE, LF, ManH and JJ analyzed the data. MarH provided *Mycoplasma mycoides* subsp. *capri* strains, ManH the proteomics infrastructure. VH, FL and JJ drafted the manuscript. All authors read and approved the final manuscript.

## Acknowledgements

We thank Helga Mogel, Sybille Schwendener, Simon Feyer and Isabelle Brodard (University of Bern, Switzerland) for their technical help. We thank Joerg Stülke for provision of the *Bacillus subtilis* strain 168 Marburg and the plasmid pHT01. We thank Patrick Viollier, Sanjay Vashee and Volker Thiel for valuable discussions.

## Supporting information

**Figure S1: Detection of transcripts of candidate TAS**

PCR using cDNA as template was used to detect transcripts. Amplicons have been separated using a 1% agarose gel. Gels on the left show amplicons obtained using gDNA as template. The gels on the right show amplicons obtained using cDNA as template and therefore presence of transcripts. Complementary DNA (cDNA) was synthesized by RT-PCR using RNA of GM12 as template. M: 1 kbp GeneRuler DNA ladder.

**Figure S2: MAUVE alignment of 16 *Mmc* genomes**

Multiple sequence alignment of different *M. mycoides* subsp. *capri* genomes using Progressive MAUVE and default parameters. Colored blocks are collinear and homologous regions. Inversions of colinear blocks are displayed below the centre line of the genome. Strain names are marked at the left of each alignment block and positions on the genomes are marked on top.

**Figure S3: Heterologous expression of different candidate toxins and antitoxins in *E. coli***

A) Immunoblot analysis of *E. coli* expressing heterologous candidate toxins and antitoxins cloned into the pBAD/His expression vector. Expression was induced by addition of arabinose. Expression was monitored over 7 h. 100 μg total protein was separated onto a 12% SDS PAGE before being transferred to a nitrocellulose membrane. PageRuler™ was used as size marker.

B) Growth curves of *E. coli* harboring candidate toxins T_752_, T_160_ or antitoxins A_753_, A_161_ or A_132_ upon induction (solid line) or repression of expression (dashed line). C) Scanning electron micrographs (magnification 10,000x) displaying *E. coli* harboring pBAD/His constructs with toxin T_133_, antitoxin A_132_ or empty vector pBAD/His either induced with arabinose or repressed with glucose.

**Figure S4: Heterologous expression of different candidate toxins and antitoxins in *B. subtilis***

A) Growth curves of *B. subtilis* harboring candidate antitoxin A_161_, toxin T_160_ or 168 Marburg with (solid line) and without (dashed line) induction of expression. B) Scanning electron micrographs (magnification 10,000x) displaying *B. subtilis* harboring pHT01 constructs with antitoxin A_161_, toxin T_160_, or empty vector pHT01 either induced with IPTG or uninduced.

**Table S1: TASmania database output file for *M. mycoides* subsp. *capri* GM12**

**Table S2: Output list of proteome analysis of *M. mycoides* subsp. *capri* GM12**

**Table S3: *M. mycoides* subsp. *capri* strains sequenced in this study**

**Table S4: TBLASTN results for candidate TAS presence in different *M. mycoides* subsp. *capri* strains and different other mycoplasmas**

**Table S5: Oligonucleotide primers used in this study**

**File S1: Plasmids constructed in this study and codon-optimized toxin and antitoxin encoding genes cloned for heterologous expression as well as TAS nucleotide sequences cloned into pMYCO1**.

